# HiC-LDNet: A general and robust deep learning framework for accurate chromatin loop detection in genome-wide contact maps

**DOI:** 10.1101/2022.01.30.478367

**Authors:** Siyuan Chen, Jiuming Wang, Xin Gao, Yu Li

## Abstract

**Motivation:** Identifying chromatin loops from genome-wide interaction matrices like Hi-C data is notoriously difficult. Such kinds of patterns can span through the genome from a hundred kilobases to thousands of kilobases. Most loop patterns are frequently related to biological functions, such as providing contacts between regulatory regions and promoters. They can also affect the cell-specific biological functions of different regulatory regions of DNA, thus leading to disease and tumorigenesis. While most statistical methods failed in the generalization to multiple cell types, recently proposed machine learning-based methods struggled when tested on sparse single-cell Hi-C (scHi-C) contact maps. We notice that there is an urgent need for an algorithm that can handle sparse scHi-C maps, and at the same time, can generate confident loop calls on regular cell lines.

**Results:** Therefore, we propose a novel deep learning-based framework for Hi-C chromatin loop detection (HiC-LDNet) and provide corresponding downstream analysis. HiC-LDNet can give relatively more accurate predictions in multiple tissue types and contact technologies. Compared to other loop calling algorithms, such as HiCCUPS, Peakachu, and Chromosight, HiC-LDNet recovers a higher number of loop calls in multiple experimental platforms (Hi-C, ChIA-PET, DNA-SPRITE, and HiChIP), and achieves higher confidence scores in multiple cell types (Human GM12878, K562, HAP1, and H1-hESC). For example, in genome-wide loop detection on the human GM12878 cell line, HiC-LDNet successfully recovered 82.5% of loops within only 5 pixels of 10k bp resolution. Furthermore, in the sparse scHi-C ODC tissue, HiC-LDNet achieves superior performance by recovering 93.5% of ground truth loops with high confidence scores, compared with that of Peakachu (31.5%), Chromosight(69.6%), and HiCCUPS(9.5%). Therefore, our method is a robust and general pipeline for genome-wide chromatin loop detection for both bulk Hi-C and scHi-C data.

**Availability:** https://github.com/frankchen121212/HiC-LDNet

## 1 Introduction

Chromatin loops occur when stretches of genomics sequences that lie on the same chromosome are in close physical proximity to each other. Multiple experimental approaches have been introduced to determine the gene loci of the looped structure. Derived from Chromosome Conformation Capture (3C) [1] technology, Hi-C [2] utilizes high-throughput sequencing technology to study the relationship between the entire chromatin DNA in the whole genome in space, and obtains high resolution rate information about the three-dimensional structure of chromatin.

Other experimental methods also provide solid genome-wide chromatin interaction information. For example, Chromatin Interaction Analysis by Paired-End Tag Sequencing (ChIA-PET) [3] incorporates chromatin immunoprecipitation (ChIP)-based enrichment to measure different aspects of chromatin organization. Similar to CHIAPET, Proximity Ligation-Assisted ChIP-seq (PLAC-seq)[4] reduces the number of input materials and improves the sensitivity and robustness by conducting proximity ligation prior to chromatin shearing and immunoprecipitation. HiChIP [5] is another approach that enables genome-wide mapping of protein-directed topological features, which lowers the input requirement over 100-fold compared with ChIA-PET. SPRITE [6] enables genome-wide mapping of higher-order interactions in the nucleus, and can detect multiple DNA and RNA molecules that interact at the same time, and get richer data than Hi-C. As a new approach to identify higher-order genomic structure, DNA SPRITE[6] is a special kind of SPRITE experiment that only assays interactions between regions of genomic DNA. STORM [7] and HiFISH [8] are two methods that utilize super-resolution imaging to reveal distinct chromatin structures, in which STORM can even differ epigenomic states in single mammalian cells.

According to previous research [9], some of the mammals’ CTCF binding protein is mostly enriched at loop anchors, while others are closely associated with enhancer-promoter interactions that can be bound by Mediator and cohesion complex. Since loop extrusion models can be used to explain the formation of topologically associating domains (TAD). Based on previous research [10], TADs are mostly formed by the ring-shaped protein complex, great proportion of diseases are closely related to TAD-like structural variations[9, 11]. For example, the duplication of TAD boundary could lead to colorectal cancer, and the fusion TADs are probable causes for Adult-onset demyelinating and leukodystrophy [9].

Although such clear structural patterns can be visually identified in Hi-C maps, it is still time-consuming and inefficient. Therefore, researchers have been working on developing different computational tools for automated chromatin loop detection. As one of the most commonly used statistical methods, HiCCUPS [12, 13] is known for its fully-automated pipeline for the annotation from Hi-C data. Another statistical method, Fit-Hi-C [14], computes the confidence estimates (p-value) on contact maps to identify chromatin interactions. HiCExplorer [15] provides a comprehensive toolbox for integrative analysis of Hi-C data. Juicer [13] and HiC-Pro [16] are two other widely used tools for the downstream analysis of Hi-C data. Mango [17] is another early proposed software that specifically works on ChIA-PET data analysis. However, statistical approaches show poor robustness to different sequencing protocols and cell types, therefore leading to a weak generalization ability. Another drawback for these statistical methods is that they heavily require intensive computational resources, thus are mostly limited to a local installation, which consumes much disk and memory during operation. Also, they do not have any scoring function so they cannot show their methods’ confidence towards the predicted loop calls. Therefore, constructing a more efficient scoring function is still one of the important goals in this research topic.

Recently, researchers have been working on applying more advanced supervised learning methods to give better detection results in variant chromosome contact maps. In Peakachu [18], a machine-learning-based framework is proposed for genome-wide interaction detection. Such a top-down method utilizes the random forest model to train a binary classification classifier and detect each sub-window across the whole genome. Being able to detect chromatin interactions on multiple cell types, such top-down-based methods needs an accurate predictor and a reliable scoring algorithm. However, it would generate multiple loop calls around the same ground truth loop, therefore a trustworthy aggregation algorithm is critical for this kind of approach. The lately proposed method, namely Chromosight [19], takes advantage of kernel convolution in computer vision to generate relatively accurate predictions from sparse matrices. It can detect multiple patterns in chromosome contact maps, including loops, borders, hairpins. To date, Peakachu and Chromosight are two of the most state-of-the-art techniques for genomewide chromatin loop detection. Nevertheless, these methods show comparative poor performance on scHi-C data, which bears high sparsity.

As single-cell analysis has become a trending topic in the field, single-cell sequencing methods target individual cells, and may help uncover many longstanding questions such as cell lineage relationship and function of DNA or histone modification at an individual-cell level [20]. With the rise of single-cell sequencing technologies, there has been an increasing demand for tools suitable for analyzing scHi-C data. Higashi [21] first utilizes a trained neural network to generate embedding for scHi-C data and impute the sparse scHi-C contact maps.

SNIPER [22] proposed a novel denoising auto-encoder for dense Hi-C reconstruction from sparse Hi-C matrices. scHiCluster [23] focuses on different patterns such as topologically associating domains (TAD) structures, and their contribution lies in the accurate clustering of different cell types from single-cell Hi-C data.

However, existing analysis tools such as scHiCExplorer [24], in most situations, cannot be used for bulk Hi-C data analysis. Therefore, it urges the need for a method that is able to possess the ability to analyze both bulk Hi-C data as well as scHi-C ones with the same tool.

Recently, deep learning has demonstrated high prediction accuracy in computer vision tasks [25] as well as in computational biology [26, 27, 28]. A simple neural network can obtain very complex underlying features, which is especially suitable for large-scale data sets and sparse dimension cases. Also, a robust deep neural network can show good generalization ability when applied to a completely new data set without losing precision and accuracy.

To this end, we propose HiC-LDNet, a novel end-to-end deep learning framework for genome-wide loop detection, which addresses topical challenges presented above:

i. HiC-LDNet can outperform existing computational methods in genome-wide loop detection throughout samples collected from different tissue types (GM12878, K562, HAP1, and H1-hESC) among multiple experimental methods (Hi-C, HiChIP, DNA SPRITE, and ChIA-PET).
ii. Our framework shows strong robustness when scanning through the extremely sparse single-cell Hi-C (scHi-C) data, and can recover the majority of the labeled loops. Compared with three state-of-the-art methods, HiC-LDNet achieves superior performance by recovering 93.5% of ground truth loops within 50kbp shift compared with that of Peakachu (31.5%), Chromosight (69.6%), and HiCCUPS (9.5%).
iii. The proposed deep learning framework can be trained in an end-to-end scheme without high memory usage and time complexity, and gives relative higher confidence scores with fewer false positive detection.

## 2 Materials and Methods

### 2.1 Overview of HiC-LDNet

As described in Fig. 1, we first train HiC-LDNet based on contact maps from different experimental methods, such as Hi-C, scHi-C, HiChIP, DNA SPRITE, and ChIA-PET. The contact map can either be read in hic format or mcool format.

**Figure 1:**
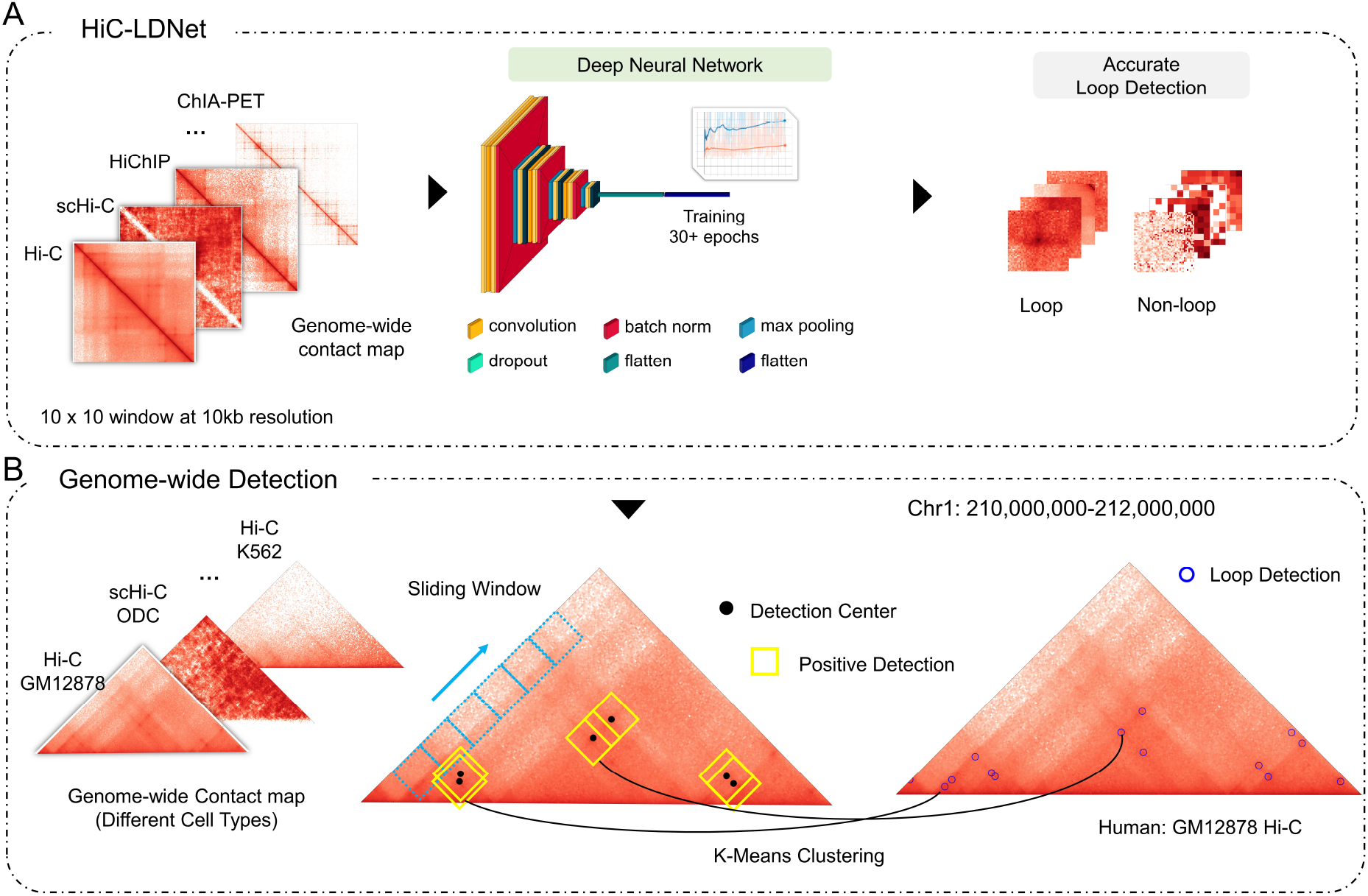
An overview of HiC-LDNet, which is applicable on multiple cell types and scHi-C. (A): After training with multiple experimental techniques, including (Hi-C, scHi-C, HiChIP, ChIA-PET and DNA SPRITE), our framework could accurately identify contact map slices with loop and non-loop classification. (B): The binary classification model is further applied on the whole genome for genome-wide loop detection with data from multiple tissue types and platforms.

#### 2.1.1 An End-to-End Trained Classifier

For positive loop classification, we extract feature representations for each loop with multiple convolutional layers with different kernel sizes. Detailed feature information can be obtained with different strides to capture the grand and local features of the given contact map (Fig. 1 A).

Since the number of positive loop samples collected in genome-wide contact maps is relatively small compared with non-loop samples. In order to imitate such imbalanced distributions, we use three times the number of negative samples compared to the number of positive samples to train our model. Specifically, the dropout layer and the batch norm layer in the network are to prevent the network from overfitting. After the model is trained for accurate loop and non-loop classification, we apply HiC-LDNet on the entire genome for genome-wide loop detection.

#### 2.1.2 Genome-wide Detection with K-means Clustering

In the prediction stage (Fig. 1 B), a sliding window algorithm is conducted to sample candidate loop window with 10 × 10 sizes of 10kb resolution, and then fed into the previous neural network. As described in Fig. 2, there will be thousands of positive candidates generated across the genome, which will definitely have overlap with each other. We further aggregate the positive detection and pool them into multiple clusters. After having the centers for all the positive detection boxes, we conducted K-means algorithm [29] to find the center of each cluster as the final loop call. The users can either choose a shorter step size for a more confident loop prediction, which may lead to a slower detection speed, or they can scan through the genome with a bigger step size in order to get a faster detection speed.

**Figure 2:**
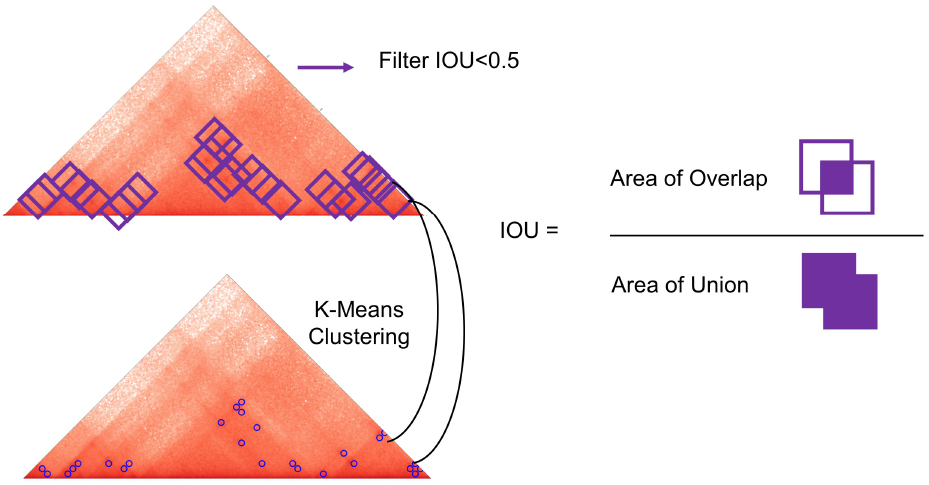
The IOU filter with k-Means clustering algorithm for loop aggregation.

When having *n* positive boxes with high soft-max probability, we initialize the clustering centers *c* = *c*_1_, *c*_2_,…*c_n_*. The task of the aggregation algorithm is to reduce the loop centers and thus reduce the false positive detection. Then, we calculate the area of Intersection over Union (IoU) between positive boxes with overlap.

When having an IoU larger than 0.5, we consider that the two boxes are detecting the same loop. After the filtering, we would define *k* loop centers and initialize *k* clustering centers. Based on the fact that the clusters are approximately Gaussian, K-means aggregation is a very proper algorithm when dealing with genome-wide detection regardless of overlap.

### 2.2 Data Set Construction

In order to prove the generalization ability of our model, we have bench-marked our model intensively across different data-sets (statistics shown in Table 1), which can be accessed through public available platforms such as 4DN Portal [30, 31, 12] and ENCODE Project [32]. In the experiments, HiC-LDNet has demonstrated a stable performance on data-sets across different tissue types as well as contact technologies. Furthermore, it has shown a significant advantage over other current detection methods when being applied on scHi-C data, whose inherent sparsity imposes a great challenge to the versatility of the detection model.

**Table 1:** The statistics of the benchmark contact maps of multiple data sets across different cell types (GM12878, K562, HAP1, H1-hESC) and multiple sequencing methods (Hi-C, ChIA-PET, DNA-SPRITE, and HiChIP). A relatively smaller number of loops are detected in scHi-C ODC cell lines. For different sequencing methods, we fix the cell line to be GM12878 and apply the same ground truth file across different sequencing protocols.

| Experimental Protocol | Cell Types | Training Loops | Distance Distribution (kbp) |  |  |  |
| --- | --- | --- | --- | --- | --- | --- |
|  |  |  | <200 | 200-500 | 500-1000 | ≥ 1000 |
| Hi-C | GM12878 | 9,166 | 3,412 | 3,402 | 1,695 | 657 |
| Hi-C | K562 | 24,475 | 15,486 | 6,665 | 1,774 | 550 |
| Hi-C | HAP1 | 8,017 | 2,601 | 3,135 | 1,643 | 638 |
| Hi-C | H1-hESC | 26,549 | 17,608 | 7,206 | 1,524 | 211 |
| Hi-C | GM12878 | 9,166 | 3,412 | 3,402 | 1,695 | 657 |
| ChIA-PET | GM12878 | 9,166 | 3,412 | 3,402 | 1,695 | 657 |
| DNA-SPRITE | GM12878 | 9,166 | 3,412 | 3,402 | 1,695 | 657 |
| HiChIP | GM12878 | 9,166 | 3,412 | 3,402 | 1,695 | 657 |
| scHi-C | ODC | 6,249 | 2,975 | 2,396 | 703 | 175 |

Each of the data set in this project is comprised of two parts, one file containing the Hi-C contact matrix, which incorporates the interaction frequency between any pairs of genome loci on any chromosome, and one file containing the coordinates of long-range chromatin interactions acquired by quality thresholding with the HICCUPS processing pipeline [12, 13], which is used to sample the positive training data for the deep learning model and as the ground truth label in later experiments.

#### 2.2.1 Data set across different tissue types

As the genes are selectively expressed among different cell types to produce cells of various functions, the chromatin exhibits cell-type-dependent three-dimensional structures, such as chromatin loops and topologically associating domains (TAD). Therefore, in order to test the versatility of our model, we have performed comprehensive experiments on our model across different cell types.

In this project, we have mainly adopted Hi-C data from four different cell lines, namely GM12878, K562, HAP1, and H1-hESC. These cell lines are chosen for their representation of the variety of human cell types. GM12878 [12] is a lymphoblastoid cell line, which has been widely used as the baseline for studying the chromatin spatial structures, DNA modification as well as genetic traits analysis. Five million cells from the GM12878 cell line are used as the input with up to nine isogenic replicates and eighteen technical replicates. K562 is the first leukemia cell line established based on human bone marrow, whose growth is regulated by cell differentiation and apoptosis. As a result, the careful investigation into the role of chromatin loop in the mechanism of gene regulation in K562 cell lines may provide us with valuable insights into the development of cancer drugs [33]. HAP1 is a nearly haploid cell line derived from cancerous cells. Since it is nearly haploid, mutations can be more easily screened and identified than common diploid cell lines. Thus, studying the loop regions in HAP1 can demonstrate more information on the relationship between chromatin structure and gene function. We have also acquired a Hi-C contact map for H1-hESC (H1 human embryonic stem) cells derived from the inner cell mass of a blastocyst, an early-stage pre-implantation embryo.

#### 2.2.2 Data set across different sequencing technologies

The Hi-C data used in this project, which measures the interaction frequencies, can be obtained via different sequencing approaches including Hi-C and its variants. In this project, we have selected Hi-C data on the GM12878 cell line acquired by Hi-C, ChIA-PET, DNA-SPRITE, and HiChIP. As each of those sequencing approaches demonstrates distinct distributions of interaction frequency and the result of each of them varies in terms of sparsity and ground truth loop counts, a generalized model should be able to exhibit equally satisfying performance across different sequencing methods.

We obtained the contact matrices sequenced with Hi-C from the ENCODE [32] project, with ChIA-PET and DNA-SPRITE from 4DN Portal [30, 31, 12] and with HiChIP from the GEO Database (GSE80820). ChIA-PET generates the contact matrix with two biological replicates and one technical replicate targeting at CTCF proteins. DNA-SPRITE generates the contact matrix with one biological replicate and seven technical replicates and is able to capture long-range chromatin interactions across a larger genome region, including inter-chromosomal interactions [6]. HiChIP uses 25 million cells from the GM12878 cell line as input with two biological replicates and two technical replicates to generate the contact matrix, and it presents a protein-centric way of sequencing interaction frequencies and shows a higher number of informative reads compared with Hi-C by 10-fold [5].

#### 2.2.3 Single Cell Hi-C Data

Besides the traditional bulk Hi-C data, single-cell technology has also led to an thriving field. The bulk Hi-C sequencing approaches may only reveal chromatin conformation at a cell population level. Thus, it is impossible to identify whether a loop or TAD appears in a specific individual cell. In comparison, single-cell Hi-C experiments can demonstrate the epigenomic state of individual cells, such as spatial interaction and orientation of chromosomes [20]. However, since the single-cell data are acquired by sampling from a cell population whose size is much more limited compared with bulk Hi-C, it imposes a great challenge to the robustness of the model used for analysis.

To address the aforementioned problem of data sparsity, we applied the SnapHiC [34] pipeline as a preprocessing tool to impute the raw single-cell Hi-C data. In specific, it imputes the contact frequencies in the original contact matrix with a random walk with start (RWW) algorithm, followed by a normalization based on linear genomic distances. In order to validate the robustness of our model on scHi-C data, we have also constructed a data set containing only scHi-C data. We obtained the single-cell sequencing data on oligodendrocytes from the dbGaP website, under phs001373.v2.p2.

### 2.3 Data Augmentation

Before feeding the data to the deep neural network, we first apply various data augmentation techniques to the training data sets [35, 36], including flipping, centered cropping, and adding random noise. During every epoch in the training process, the samples are flipped along their main diagonals with a probability of 0.5, while the original data set remains unmodified. Furthermore, the samples are cropped at the center. While detecting loop regions with a sliding window, a chromatin loop might not always occur at the center of the window. Therefore, centered cropping can help boost the model’s detection performance. Moreover, random Gaussian noise is applied to all the training samples to maintain the generalization ability of the model. Experiments demonstrate that data augmentation addresses the problem of positive data scarcity as well as the class imbalance issue. In addition, it helps our model avoid overfitting and obtain better generalization power.

### 2.4 Imbalanced Training and Data Quality control

To prepare our framework for the genome-wide detection, we train the neural net based on imbalanced samples, with 9,119 loop samples and 27,351 non-loop samples for Hi-C GM12878 cell line, 23,928 loops and 71,784 non-loop samples for Hi-C K562 cell line, 26,521 loops and 79,563 non-loops for H1-hESC cell line, and 7,963 loops and 23,389 non-loops for HAP1 cell. For each cell type, we have randomly generated negative samples across the genome with low IOU across the loop regions.

Since the majority region in Hi-C matrix is incomplete (missing some attribute values of interest), the matrix values from different experimental protocols are also variant in range. We conduct quality control over different cell types as well as Hi-C matrices across multiple experimental protocols. We first log-transform the matrix slices, and then each window is normalized to the range (0,1) to unit variance and zero means. Such implementation enables us to improve the data quality and prevent gradient explosion during training and accelerate the convergence of our model.

### 2.5 LDAM Loss for Imbalanced Training

The training set sample sizes for each category are not balanced. Meanwhile, the genome-wide testing standard requires good generalization ability for each category and is even more concerned about the performance of the minority category [37, 38], which in our case is the rare chromatin loops. When we train the model on such an imbalanced dataset, standard Cross-Entropy Loss (CE) fails to optimize our network with high precision.

By giving a larger margin to the minority loop class, we need to shift the actual classification boundary so as to reduce the difficulty of classifying the minority class. Label-Distribution-Aware Margin (LDAM) loss [39, 40] proposed a class-dependent margin for the multiple-class classification. The margin Δ*_j_* for the *j*-th class is defined by the number of class samples to give them an appropriate shift:

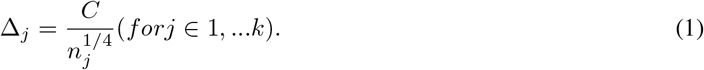

Here, *k* is 2 in our binary classification setting, and *C* is a hyper-parameter to be tuned. In our training, to minimize marginal generalization boundary. The loss is defined by calculating the probability that the logit of the actual label *y* is less than the other label.

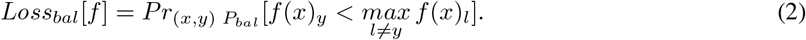

Therefore, the final loss after Softmax activation for imbalanced training can be written in

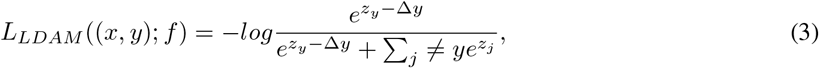

where Δ is a hyperparameter defined by the number of sample sizes in Equation 1.

After systematical experiments, we find that LDAM loss shows significant improvement compared with Cross-Entropy Loss (CE) and Focal loss. Therefore, when optimizing our prediction model, we use the LDAM loss to ensure a better decision boundary and a more robust performance accuracy on genome-wide detection.

### 2.6 Evaluation Metrics for Training

Essentially, the deep neural network is trained as a binary classifier and then applied on large-scale Hi-C contact matrices to detect chromatin loop regions. As a result, we use the F1-score, which is the harmonic mean of the model’s precision and recall, as the key performance metric to measure the model’s training results. In specific, the F1-score is defined as

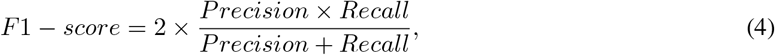

where

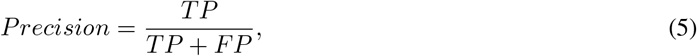

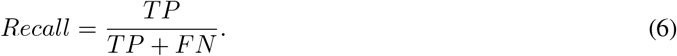

Notice that here, the accuracy of the prediction is not as informative in our experiments as F1-score since the training data set is highly imbalanced, and loops are rarely found in a genome-wide contact matrix.

### 2.7 Genome-wide Detection Evaluation

In order to validate each algorithm’s prediction performance across the genome, we calculate the distance between the nearest predicted loop and the ground truth loop caller and calculate their Euclidean distance (shift). That is, the smaller the euclidean distance (shift) from the prediction to ground truth, the better the model performs.

Therefore, the overall distance list from the loop anchor *A* and the prediction *P* for method across the entire genome can be calculated in the following formula:

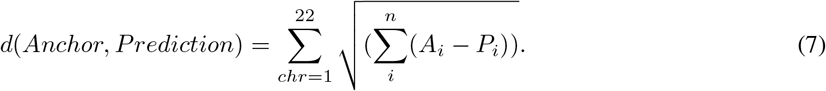

We then calculate the number of positives that falls in close range of distance from the ground truth anchor under a certain score threshold. For most loop calling algorithms, a detection is considered positive if the distance between the predicted loop call and the ground truth loop position is less than 10 times of pixel size, which in our case is 10kb resolution, thus making an accurate detection falls in 100kbp distance from the loop anchor.

### 2.8 Experimental Details

The neural network is composed of two convolutional layers, both of them are of kernel size 3 and stride 1. To overcome the vanishing gradient problem and make our models learn faster, the Rectified Linear Unit (ReLU) activation function is connected after each convolutional layer. Furthermore, to prevent gradient dispersion, we also apply batch normalization during our training. In practice, on each data set, we have trained our network with a learning rate equal to 0.001 for 25 epochs with Adam optimizer, betas=(0.5, 0.999). The dropout ratio is set to 0.1 after the first and second convolution layer to prevent over-fitting.

As for HiC-LDNet’s time complexity, it requires about 20 minutes to train on each data set. During genomewide detection, an average speed of 25 s/Mbps can be achieved for the whole process. The final output of our program consists of a bedpe file and the detection figure. The bedpe file labels all the coordinates for the detected loops with the first and fourth column indicating the chromosome, and the seventh column giving the model’s confidence score.

## 3 Results

### 3.1 HiC-LDNet accurately recover majority known interactions from Hi-C GM12878

To test our model’s viability on known Hi-C data, we first trained and tested HiC-LDNet on 9,119 known loops on the homo sapiens GM12878 cell line. As shown in Fig.3A, we can see a higher binary classification accuracy on the test set with Hi-C contact maps (95.2%) than that of ChIA-PET(74.8%), DNA SPRITE (74.1%), HiChIP (89.2%), and scHi-C (91.2%).

**Figure 3:**
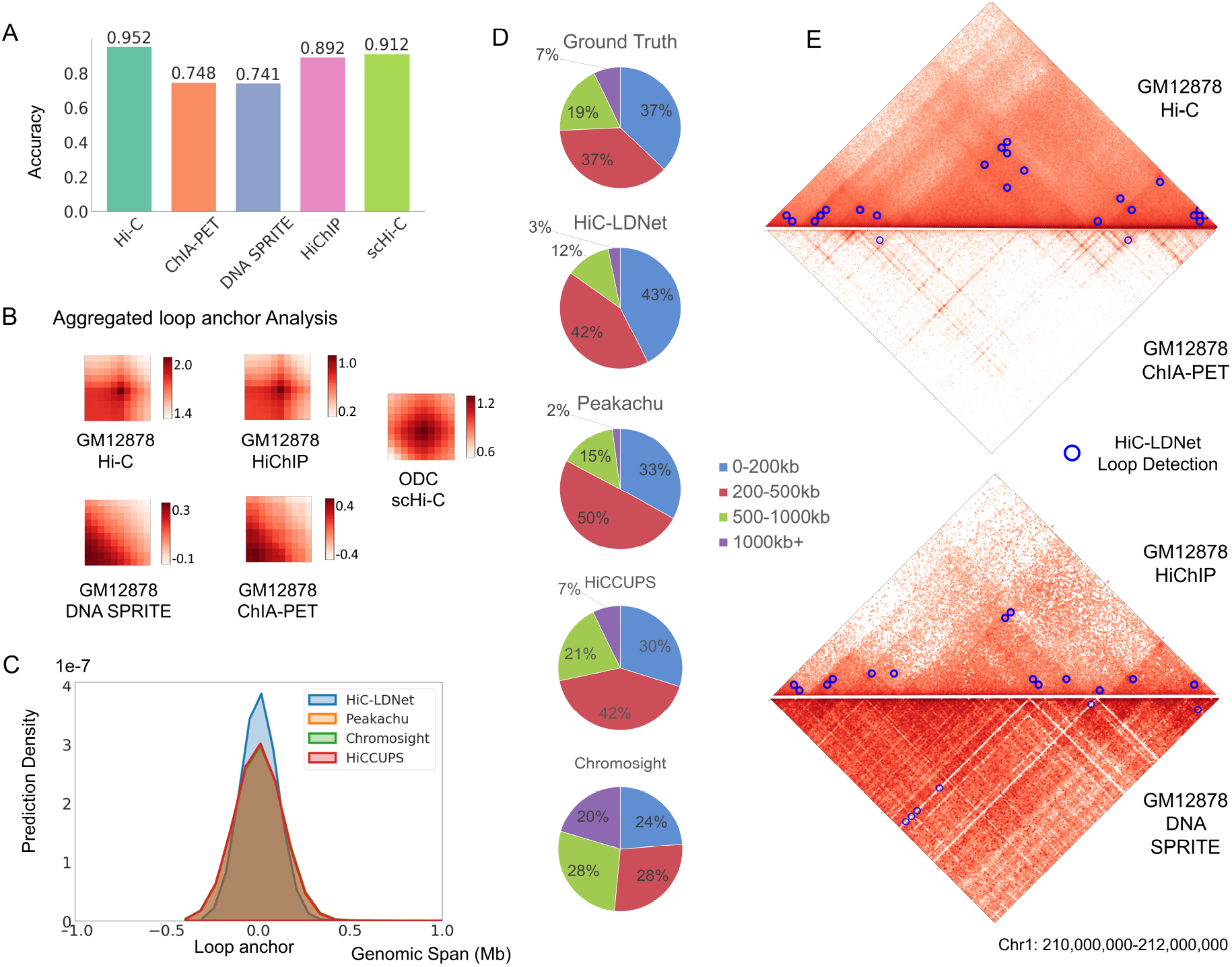
Application of HiC-LDNet on GM12878 Hi-C, as well as ChIA-PET, HiChIP, and DNA SPRITE. (A): The binary classification accuracy of HiC-LDNet’s performance over 5 different experimental protocols. (B): The aggregated loop anchor patterns for all the positive window slices with 10 × 10 pixels in 10kb resolution. (C): On the GM12878-Hi-C, the prediction density comparison of HiC-LDNet, Peakachu, Chromosight, and HiCCUPS. Hic-LDNet extracted comparatively more loop calls in close regions around the loop anchor. (D): On GM12878-Hi-C, the distance distribution of loops uniquely detected by HiC-LDNet and other baseline methods. (E): The example genome-wide loop detection result on a fraction of chromosome 1 base on our proposed HiC-LDNet on different experimental protocols, including Hi-C, ChIA-PET, HiChIP, and DNA SPRITE.

As can be seen in the aggregated peak analysis in Fig.3 B, a strong Hi-C enrichment signal emerged in the center of most ground truth loop regions. From a two-sided t-test for all the loop calls and the sub-sampled nonloop calls, a p-value of 2.864 e-08 is achieved to show the unique pattern of loop calls among the GM12878 cell line. Such a pattern is still significant when we conduct analysis on GM12878 HiChIP (p = 5.87 e-2) and on oligodendrocytes (ODC) for scHi-C data (p = 1.06 e-21). However, the pattern is not as much significant in GM12878 DNA SPRITE (p = 0.21), as well as GM12878 ChIA-PET (p = 0.54) in loop and non-loop regions.

It is worth noticing that the average log10 value of the GM12878 Hi-C (1.88) loop is relatively higher than that from HiChIP (0.89), DNA SPRITE (0.29), and CHIA-PET (0.96). Therefore, it demonstrated a clearer loop pattern in Hi-C matrix other than other experimental protocols.

In Fig 3 D, we see a majority (37%) of ground-truth interactions clustered in a short range (¡200kb), and another big portion (37%) in medium range (200-500kb). Our proposed HiC-LDNet gives a prediction that reveals such percentages, with (43%) loop calls in (¡200kb) range and (42%) within 200-500kb range. The distribution of loop patterns in the prediction is consistent with the training set patterns for our method.

### 3.2 Comparison of HiC-LDNet with Peakachu, Chromosight, and HiCCUPS on GM12878 Hi-C

To benchmark the performance of HiC-LDNet, we compare it with 3 lately released methods. One is the Random-Forest based loop detection methods Peakachu[18]. Another is the loop caller from Chromosight [19], which is a kernel-convolution-based method inspired by computer vision. The last one is HiCCUPS [12], which is a statistical method that utilizes the Poisson test with modified Benjamini-Hochberg adjustment to determine loop interactions.

Statistically, loop calls generated by HiC-LDNet are much closer to the loop anchor compared with the other loop calling algorithms. As can be seen from the kernel density estimate (KDE) plot in Fig.3C, the density of loops clustering around the loop anchor is much higher in our proposed method. We have also observed the LDNet’s prediction patterns to be highly correlated with the ground truth loop size distribution.

We systematically compare the loop uniquely detected by each method on the entire 22 regular chromosomes from the human GM12878 cell line. Under the same loop call threshold, 91.6% of the loops are accurately detected within only a 100kbp shift from the ground truth label, of which 82.5% of them are detected within only 50kbp shift, compared with that of 80.0% of Peakachu and 82.0% of HiCCUPS. Chromosight shows a very good performance of recovering 90.2% of the ground truth loops within a small shift, while it gives a number of missed detection with low confidence scores. For example, only 1.1% of HiC-LDNet’s predictions are far away from the ground truth (¿1000kb), compared with 1.3% of Peakachu, 1.2% of Chromosight, and 1.4% of HiCCUPs.

### 3.3 Generalization ability of HiC-LDNet on different cell types and experimental methods

Besides Hi-C, our framework can be further applied to contact data of multiple experimental protocols. We have acquired publicly available cell line data sets from multiple platforms, including human GM12878 cell lines of Hi-C, ChIA-PET, HiChIP, and DNA SPRITE. We have also conducted experiments on multiple human cell lines (GM12878, K562, HAP1, and H1-hESC).

In the GM12878 cell line, multiple contact maps are tested for genome-wide detection results. In Fig. 3 E, we demonstrate HiC-LDNet’s prediction performance under different experimental protocols. Rare loop calls are detected in CHIA-PET (1,239 out of 9,126) and DNA SPRITE (584 out of 9,126) contact maps due to a high signal-to-noise ratio, while our method achieved a promising prediction performance on HiChIP contact maps. Also, in Fig. 4, HiC-LDNet shows its liability of accurately calling most of the loops on genome-wide detection.

**Figure 4:**
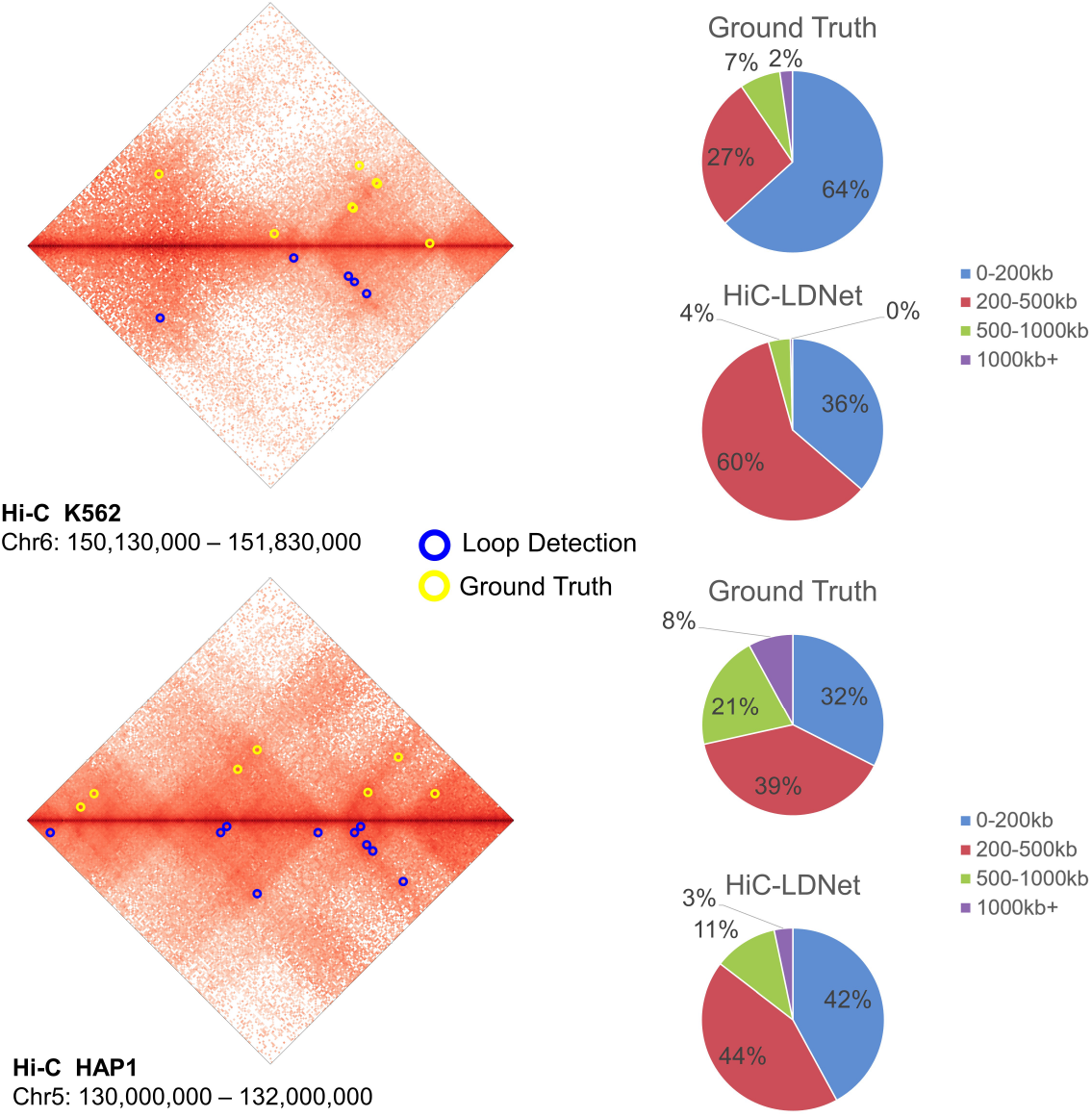
Example of HiC-LDNet’s genome-wide prediction result on chromosome 6 (150,130,000-151,830,000) of K562 cell line and on chromosome 5 (130,000,000-132,000,000) of HAP1 cell line. On the right panel, we see that HiC-LDNet’s detected loop distance distribution is highly correlated to the ground truth distribution.

Apart from that, it is worth noticing in the loop distance pie chart Fig. 4, HiC-LDNet demonstrates its strong ability to uniquely label loop calls with low distance. In K562 cell lines, a total number of 24,475 chromatin loops are labeled according to the ground truth, HiC-LDNet is able to provide 28,195 predictions, with the majority detection loops (36%) clustered within 200kb, and a few larger than 100kb. Similarly, in HAP1 cell line, a total number of 8,017 loops were labeled and HiC-LDNet managed to give 5,611 confident predictions with scores higher than 0.9, and 12,969 predictions with scores higher than 0.8.

### 3.4 HiC-LDNet shows strong robustness applying on sparse scHi-C Data compared with Peakachu, Chromosight, and HiCCUPS

Single-cell Hi-C (scHi-C) can identify cell-to-cell variability of three-dimensional (3D) chromatin organization, but the sparseness of measured interactions poses an analysis challenge. HiC-LDNet can perfectly deal with such challenges and show strong robustness in sparse scHi-C contact matrices. From the comparison with our proposed method’s loop detection in the visualized scHi-C map in Fig. 5D, HiC-LDNet can accurately identify multiple loops with few false-positive predictions compared with Chromosight and Peakachu. Statistically, 93.5% (5,675 out of 6,073) of HiC-LDNet’s predictions falls into 50kbp or less shift from the ground truth loop anchor, compared with 31.5% (1,911 out of 6,073) of Peakachu, 69.6% (4,225 out of 6,073) of Chromosight, and 9.5% (575 out of 6,073) of HiCCUPS. It is worth noticing that, like other cell types, we have also acquired scHi-C matrix with 10kb resolution. Therefore, it indicates that most predictions from our framework have within 5 pixels shift from the ground truth label.

**Figure 5:**
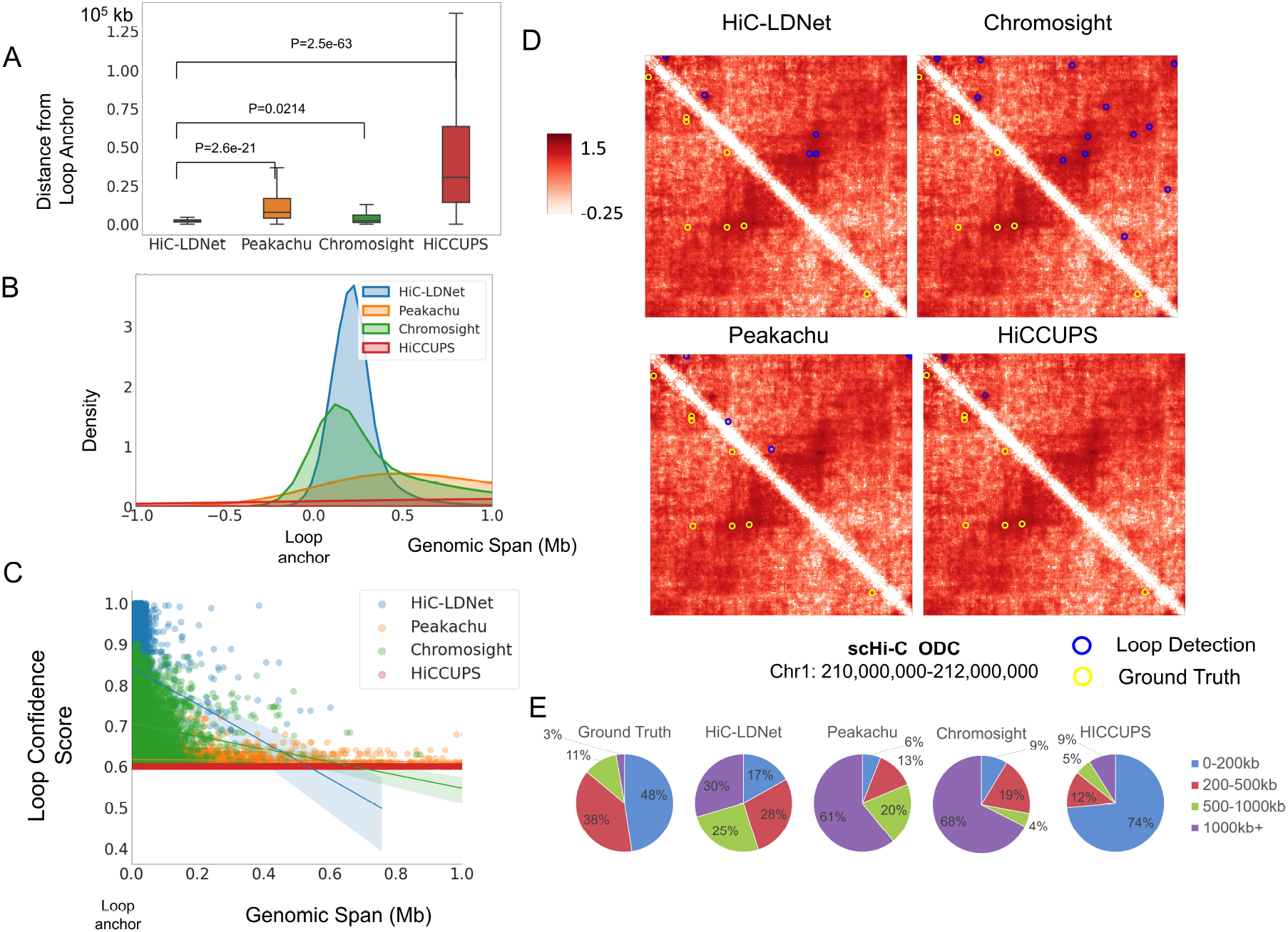
Comparison of HiC-LDNet with other genome-wide loop calling algorithms performance on the scHi-C ODC cell line. (A): A box plot showing the distance distribution from the detected loop to the loop anchor with a p-value calculated from a 2-sided t-test between each distribution. **(B):** The kernel density estimation (KDE) of the genomics span shows where the predicted points are clustered. Most of HiC-LDNet’s predictions are clustered around the loop anchor. (C): The distribution of the highly scored loop calls with respect to their position among the 4 algorithms. HiC-LDNet shows a strong confidence score compared with the state-of-the-art algorithms. (D): Example of the predicted results of 4 loop calling algorithms on the chromosome 1 chromatin interaction for the scHi-C ODC cell line. (E): The distribution of loop distances of the ground truth and each algorithm’s prediction.

To further validate our framework’s robustness on the scHi-C oligodendrocytes (ODC) data, we first draw the box plot of the distance from the loop anchor for 4 model’s prediction loop calls. As can be seen in Fig 5A, HiC-LDNet has the majority number of prediction loops clustered within a short distance from the ground truth peaks. From a two-sided test that shows the significant difference in the shift distance, HiC-LDNet showes significant improvement from Peakachu (p=2.6e-21), Chromosight (p=0.0214), and HiCCUPS (p=2.5e-63).

Another experiment is to see how confident each model is based on its prediction scores. When each loop calling algorithm gives a score for a detected loop (score is usually a value between 0 to 1), it demonstrates the confidence level of the model’s prediction towards that loop call. For example, Peakachu [18] applies a greedy pooling algorithm and gives the best-scored contacts from high-probability pixels. In our method, we have averaged the loop scores within overlapped positive detections from the Softmax activation function. As can be seen in Fig5 C, the majority of the loop scores are centered above 0.9, compared with that of Chromosight and Peakachu. Since HiCCUPS [12] is a statistical method with no confidence score, we have simply labeled all its prediction loops according to the threshold.

We have further analyzed the overlap between each methods’ peak calls. As can be seen in Fig.5 E, HiC-LDNet identified a unique set of chromatin interactions from Peakachu (52,859) and Chromosight (21,588). While most of the ground truth loops for scHi-C ODC are basically clustered within 200kb regions. Rarely has Peakachu or Chromosight managed to detect those regions and give a high confidence score. However, our proposed method shows an average split between the loop distance from (0-200kb) and (200-500kb) in high confidence scores.

## 4 Discussion and Conclusion

Facilitating genome-wide loop detection is a crucial part of understanding how these structures regulate numerous cellular processes, including but not restricted to replication, termination, and chromosome segregation. Therefore, an accurate and robust loop calling algorithm is important for revealing such processes and studying gene regulation. In this work, we propose a novel deep-learning-based framework to accurately detect chromatin interaction loops in variant genome-wide contact maps. We use the top-down scheme to train our neural networks, and our framework is able to eliminate the effect brought by the sparsity of scHi-C data. We analyzed different loop regions of variant human cell lines, including GM12878, K562, HAP1, and H1-hESC.

Compared with the existing state-of-the-art methods, the evaluation of our program demonstrated that HiC-LDNet significantly outperforms those algorithms and is more robust to scHi-C data with huge sparsity. Considering the time complexity of our method, HiC-LDNet could finish its prediction at an average 25 s/Mbp speed across the entire genome at 10kb resolution. Though not the fastest method, HiC-LDNet still achieves competitive speed compared with the state-of-the-art methods.

To further improve HiC-LDNet, future works remain to be challenging and promising. For instance, with the advantage of HiC-LDNet, it can be further applied on scHi-C matrices for feature extraction, hypergraph embedding, genome structure characterization, and cell-type classification. Furthermore, with both TADs and sub-TADs are populated by focal points of interaction (loops), HiC-LDNet can be further applied for TAD identification with a decrease of interactions at the boundary regions. This work is a step toward this goal while we understand that this topic is open for further investigations from regular Hi-C and sparse scHi-C contact maps.

